# Identification of *Ensifer meliloti* genes required for survival during peat-based bioinoculant maturation by STM-seq

**DOI:** 10.1101/2022.09.19.508585

**Authors:** Mauricio J. Lozano, Ezequiel G. Mogro, M. Eugenia Salas, Sofía A. Erdozain, Nicolás E. Zuber, Anke Becker, Antonio Lagares

**Affiliations:** Instituto de Biotecnología y Biología Molecular — CONICET CCT-La Plata, Departamento de Ciencias Biológicas, Facultad de Ciencias Exactas, Universidad Nacional de La Plata, La Plata, Argentina; LOEWE Center for Synthetic Microbiology and Faculty of Biology, Philipps University, Marburg, Germany

**Author notes:** Corresponding author Mauricio J. Lozano. These authors contributed equally to this work (Mauricio J. Lozano, Ezequiel G. Mogro, M. Eugenia Salas).

**Keywords:** signature tagged mutagenesis, phenomics, inoculant, rhizobium, peat

## Abstract

Rhizobial inoculants are sold either as rhizobia within a liquid matrix; or as rhizobia adhered to granules composed of peat prill or finely ground peat moss. During the production of peat-based inoculants, immediately after mixing the rhizobia culture with partially dry sterile peat, the inoculant is stored for a period of 4-5 weeks, inducing a series of changes that results in an increased capability of the rhizobia to survive in the seeds. The number of viable rhizobia on preinoculated seeds at the point of sale, however, is often a limiting factor, as is the inefficiency of the inoculant bacteria to compete with the local rhizobia for the host colonization. In the present work, we used STM-seq for the genomewide screening of *Ensifer meliloti* mutants affected in the survival during the maturation of peat-based inoculant formulations. Through this approach, we identified hundreds of genes that proved to be relevant to this process. These results also provide a base knowledge that could be used to more completely understand the survival mechanisms used by rhizobia during the maturation of peat-based inoculants, as well as for the design of new inoculant formulations.

**Highlights:** Rhizobial inoculants provide an ecological means of nitrogen fertilization compatible with the implementation of sustainable agricultural practices. Their successful usage, however, suffers from two main limitations: the low number of viable rhizobia on preinoculated seeds at the point of sale, and the inefficiency to compete with the local rhizobia for host colonization. Here, we used a high-throughput mutant-screening technology, STM-seq, to uncover which rhizobial genes are involved in the rhizobial survival during the preparation and storage of peat-based inoculant formulations. Our findings provide useful information about the stresses faced by rhizobia during peat-inoculant maturation and storage, which could assist both for the selection of better rhizobial strains, and for the improvement of the inoculant formulations.

**Graphical Abstract:** 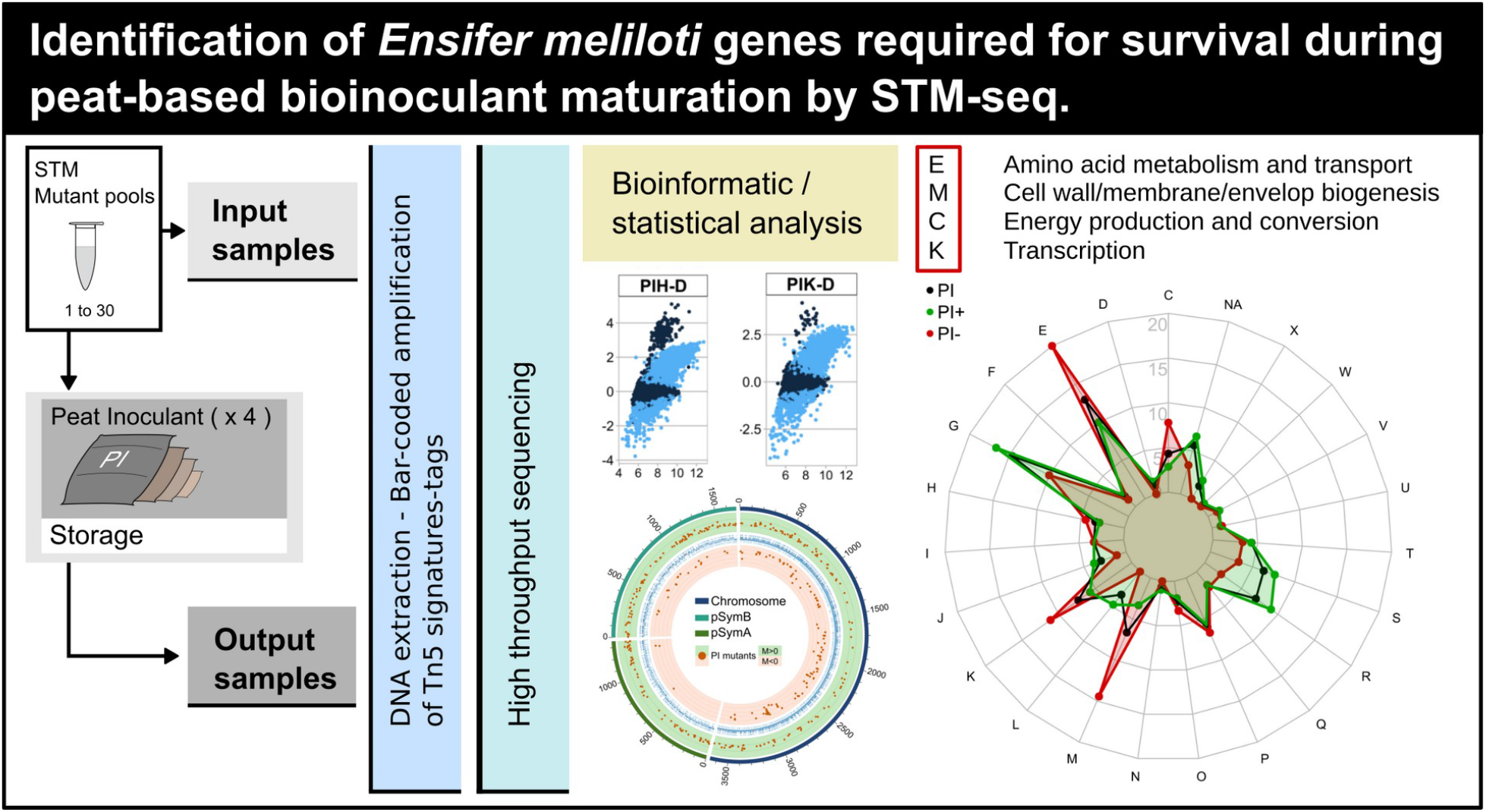

## 1. Introduction

Rhizobia are soil dwelling bacteria that interact with legume plants through a complex series of coordinated communication and differentiation events, resulting in the formation of highly specialized root structures, the nodules, in which they differentiate into N_2_-fixing bacteroids (Gourion et al., 2014; Oldroyd, 2013; Poole et al., 2018). Rhizobium-legume symbioses provide a major proportion of the fixed N_2_ in the biosphere, with the inoculation of seeds with rhizobia having become a routine practice employed since the early 20th century (Brockwell and Bottomley, 1995; Smith, 1992; Stephens and Rask, 2000). Nevertheless, the availability of cheap fertilizer N, the need for higher crop yields, and the generalized use of intensive agricultural practices, have produced a decline in the biological N_2_ fixation, even at the cost of significant environmental damage (Graham and Vance, 2000; Olivares et al., 2013). In that scenario, the use biological N_2_ fixation has proved to be a profitable approach to arrest the decline of nitrogen fertility of the soil (Peoples et al., 1995), although much room for improvement still remains. Two main limitations have been observed: the low number of viable rhizobia on preinoculated seeds at the point of sale (Catroux et al., 2001; Deaker et al., 2004; Graham and Vance, 2000) and the inefficiency to compete with the local rhizobia for the host colonization (Catroux et al., 2001; Graham and Vance, 2000; Maier and Triplett, 1996; Silva et al., 2007).

Currently, most inoculants are sold in three ways: as rhizobia within a liquid matrix that is applied to the seeds or introduced into the sowing furrows, or as rhizobia adhered either to granules composed of peat prill or to finely ground peat moss. While the granule products are placed directly in the furrow, peat moss is applied to the seeds before sowing (Deaker et al., 2004; Stephens and Rask, 2000). During the production of peat-based inoculants, immediately after mixing the rhizobia culture with partially dry sterile peat, the inoculant is stored for a period of 4–5 weeks (Smith, 1992). This storage period, commonly referred as “maturation” or “curing”, induces a series of changes that result in an increased capability of the rhizobia to survive in the seeds (Albareda et al., 2008; Materon and Weaver, 1985; Roughley and Vincent, 1967). The absence or shortening of this storage period produces drastic effects on the viability of the bacteria in the seed (Materon and Weaver, 1985). The improved survival of rhizobia in the seeds after the maturation process involves morphologic changes such as thickening of the cell wall, fewer polyhydroxybutyrate granules and periplasmic space occlusion (Dart et al., 1969; Feng et al., 2002). One of the factors affecting the survival of rhizobia in legume seeds is desiccation (Deaker et al., 2004). A better understanding of desiccation stress tolerance might help to understand in part the adaptation undergone within peat, and how that process improves survival in the legume seeds.

Only a few studies have been reported that aimed at the molecular characterization of the biological processes taking place in rhizobia during inoculant formulation, maturation and shelf-life. Casteriano *et al*. (2013) demonstrated that rhizobia treated with aqueous peat extracts constituted an informative system for studying the adaptation undergone within peat inoculants. They studied desiccation tolerance on *Rhizobium leguminosarum bv. trifolii* TA1 and *Bradyrhizobium japonicum* CB1809 and found that peat extract treated bacteria presented a better tolerance to desiccation by vacuum, increased levels of intracellular trehalose, and were similar to bacteria recovered from peat. This system also enabled the authors to perform polyacrylamide-gel electrophoresis (PAGE) based proteomic analysis of peat-extract treated bacteria and, as a result, identify 14 (TA1) and 8 (CB1809) differentially produced proteins. The most relevant proteins induced under those conditions were related mainly to stress response, translation, and transporter activity. In a more recent publication, Atieno *et al*. (2018), using two dimensional PAGE proteomics, identified differentially produced proteins on peat extracts in four rhizobial strains. Proteins exhibiting the greatest increase in abundance were those involved in amino acid and carbohydrate transport and metabolism, posttranslational modification, stress-response, and cellular-defense mechanisms. In addition, the authors found that growth in peat extract caused damage to cell membranes and exposed rhizobia to sublethal stress, resulting in the differential production of several stress-induced proteins that may be responsible for the observed peat induced protection against desiccation.

In the last decade, second generation sequencing technologies enabled the simultaneous phenotypic analysis of thousands of mutants through transposon-insertion-sequencing methods (van Opijnen and Camilli, 2013), such as Tn-seq (van Opijnen et al., 2009), TRAdis (Langridge et al., 2009), Inseq (Goodman et al., 2009), RB-Tnseq (Wetmore et al., 2015) and STM-seq (Salas et al., 2017). Although those methods were applied mostly to the characterization of pathogenic bacteria, they have recently been used for the study of soil and plant associated bacteria (Bishop et al., 2014; Bishop and Rachwal, 2014; de Moraes et al., 2017; diCenzo et al., 2018; Duong et al., 2018; Liu et al., 2018; Price et al., 2018; Royet et al., 2019; Sivakumar et al., 2019), with highly informative results regarding the identification of rhizobial genes required for growth under different conditions (Perry et al., 2016a; Wheatley et al., 2017), as well as during the interaction with the plant host (Perry et al., 2016b; Salas et al., 2017; Wheatley et al., 2020). Nevertheless, these methods have still not been applied to the identification of genes required for the survival of bacteria during inoculant manufacturing.

In the present work, we report the use of STM-seq (Salas et al., 2017) for the genomewide screening of *Ensifer meliloti* mutants affected in their survival during the maturation of peat-based inoculant formulations. Through this approach, we have identified hundreds of genes that proved to be relevant to this process, either beneficially or detrimentally. In addition, our results provide fundamental information that will be of use to more thoroughly understand the mechanisms used by rhizobia to survive during the maturation of peat-based inoculants as well as for the design of new inoculant formulations.

## 2. Materials and methods

### 2.1. Bacterial strains and growth conditions

Thirty sets of a *E. meliloti* signature-tagged mini-Tn5 mutant library (Pobigaylo et al., 2006; Serrania et al., 2017) were used. *E. meliloti* cells were grown at 28 °C in tryptone-yeast (TY) complex medium (Beringer, 1974) or in Evans defined minimal medium (Evans et al., 1970), containing glucose (10 g/L) and ammonium chloride (0.7 g/L) as the respective carbon and nitrogen sources. For solid media, 15 g of agar per liter of medium were added. When required, antibiotics were used in the following concentrations: neomycin (Nm, 120 µg/ml), streptomycin (Sm, 400 µg/ml) and cycloheximide (Cx, 100 µg/ml).

### 2.2. Screening of STM mutants in peat

For the screening experiments, we used commercial bags containing 100 g of gamma-radiation sterilized peat. Mutant sets (Pobigaylo et al., 2006) were grown in 10 ml of TY for 20 h to an optical density at 600 nm (OD_600_) of 2–5. Each liquid culture was diluted 1:8 in peat diluent solution [Sucrose (128 g/l), KH_2_PO_4_ (0,68 g/l) and K_2_HPO_4_ (0,88 g/l)]. A part of this diluted culture was stored and used as a reference [input sample, approx 1×10^9^ colony-forming units per ml (CFU/ml)]. Peat bags were inoculated by injection with 5 ml of input sample, and for each mutant set, 4 replicate bags were used. After covering the injection site with a labeled sticker, each bag was massaged manually to homogenize the content. The inoculated peat bags were stored in closed boxes for two months at 22 °C (maturation period).

After the maturation period, 5 g of inoculated peat from each bag was mixed with 10 ml of TY and vortexed for 10 s. After 5 min the supernatant was passed through filter papers (grade 1F, particle retention 5 to 6 μm), and 200 µl of the filtered liquid containing the bacteria were used to inoculate 10 ml of TY media containing Sm, Nm and Cx, followed by growth for 15 h (between 4 and 5 generations). For storage, 900 µl of culture were mixed with 100 µl of 10X lysis solution and centrifuged for 1 min at max speed, 900 µl of supernatant were discarded, and finally the pellet was resuspended in the remaining volume to constitute the output sample. Samples were processed as indicated in Salas *et al*. (2017) for DNA extraction.

### 2.3. Real-time polymerase chain reaction (qPCR) amplification of mutant signatures

The PCR amplifications of the H and K signatures with the bar-coded primers were carried out in a real-time thermal cycler qTOWER (Analityk Jena) with the Maxima SYBR Green qPCR master mix (Thermo Scientific) in a total reaction volume of 10 µl, containing each oligonucleotide at 0.6 mM and 2.5 µl of DNA template prepared as described in Salas *et al*. (2017). The cycling conditions used were as follows: an initial denaturation at 95 °C for 10 min; followed by 35 cycles of 10 s at 95 °C for denaturation, 5 s at 65 °C for annealing, and 15 s at 72 °C for elongation. Table S1 lists all 76 different primers used in the present work.

### 2.4. Mixes of PCR products, high-throughput DNA sequencing, and sequence data processing

In order to obtain a similar number of reads for each experimental condition, equivalent amounts of each one of the PCR amplification products were mixed. To that end, the maximum intensity achieved in the qPCR reaction—i. e., the fluorescence intensity obtained for each tube in the plateau—was taken into consideration. In order to mix comparable amounts of the PCR products, volumes between 2.5 and 4.5 µl of each sample were mixed.

### 2.5. DNA sequencing and bioinformatic analysis

Paired-end sequencing of the qPCR products was performed by means of the Illumina HiSeq 1500 platform at the CeBiTec (Center for Biotechnology), Bielefeld, Germany. As a result, a total of 2.58 × 10^8^ assembled paired-end reads was obtained as two 33 Gb FastQ files (one file per Illumina-sequence lane). The sequence quality was evaluated by means of the FastQc software (Andrews, 2010). Both FastQ files were concatenated, converted to fasta format, and size-filtered to select for sequences with sizes longer than 110 bp through the use of a local instance of the Galaxy platform (Blankenberg et al., 2010). After this procedure a total of 2.36 × 10^8^ sequence reads matching this criterion was obtained. The representation of each mutant in a given mix was established through the count of their corresponding signatures. The identification and quantification (count) of each mutant—*i. e*., a combination of the H and K signatures in a given mutant mix, as provided by the P2 and P4 bar-codes—under each experimental condition—*i*.*e*., peat formulation, and replica; provided by the P1 and P3 bar-codes (Salas *et al*., 2017)—were performed with Perl scripts written for this purpose (https://github.com/maurijlozano/STM-seq-count). The resulting quantitative input and output data for each mutant under each experimental condition were finally consolidated by means of a standard relational database management system. This process was independently applied to the H and K signatures in order to evaluate the concordance (robustness) of the experimental procedure from the qPCR step to the quantitative analysis, which results provided additional statistical support for the entire experimental procedure. A total of 1.08 × 10^8^ reads were analyzed, where Table S2 lists the detailed data for each condition and replicate.

### 2.6. Data analysis

The consolidated data for each experimental condition (two biological conditions with four biological replicates each) and each of the 30 STM mutant mixes consisted in a 412 × 8 raw count matrix (for either the H or the K data, and for each mutant mix) including: 412 rows for the different mutants analyzed, four columns for the input replicates, and four columns for the output replicates. For each of those conditions, the raw read count associated with each mutant was normalized by the total read count of the sample. Then the M value [log fold change = log_2_ (Output reads / input reads)] and A value [½ log_2_ (Output reads * input reads)] were calculated. A pseudosum of 1 was applied in order to avoid infinite values. MA plots were constructed indicating that further normalization of the data was required. Two main biases were observed: (1) The average M-value was smaller than 0, and (2) a correlation between the M and A values was observed. To improve the data analysis, each of the count matrices was analyzed by means of the TCC R package (Sun et al., 2013). Two alternative pipelines were followed, edgeR coupled with iDEGES/edgeR normalization, and DESeq2 coupled with iDEGES/DESeq2 normalization. The results of both pipelines were consolidated for the selection of differentially expressed genes (DEG). Both pipelines were run with a false discovery rate (FDR) set to 0.01 and a floor-PDEG of 0.005 for normalization, and with a FDR of 0.005 for differential expression estimation. In addition, the low count filtering was set at 300. The results corresponding to H and K signatures were analyzed independently (Table S2), and presented a very good correlation (Table S3), especially in accounting for the DEGs. In addition to the DEG estimation by the TCC software, we considered the DEG genes to be those with M-values smaller than −1 or larger than 2. This criterion was defined on the basis of the volcano and MA plots, and the small bias towards positive fitness mutants in the results (Table S3).

### 2.7. Validation assays

In order to validate the phenotypic results obtained by STM, the survival in peat inoculant (PI) of a set of selected mutants was individually assessed. To that end, sterile peat was inoculated with *ca*. 10^8^ CFU of either wild-type *E. meliloti* 2011 or a given mutant onto 1.5-ml polypropylene tubes. These samples were incubated for 15 days at 10 °C. The bacteria from the peat treatments were recovered at times 0 and 15 days thereafter by adding 500 µl of TY broth to the initial inoculate, mixing with a vortex, and taking 100 µl of the supernatant after the sedimentation of the peat. The recovered bacteria were then diluted and plated in TY agar plates supplemented with Sm in order to count the viable cells that had survived the treatment. The CFU for the inoculum and the treated samples were counted, and the proportion of surviving bacteria was calculated (Table S4).

## 3. Results

### 3.1. Genome-wide identification of E. meliloti mutants with altered fitness in peat-based inoculant formulations

In order to identify *E. meliloti* genetic determinants associated with an altered fitness in peat based inoculant formulations, we used a Tn5-STM approach combined with high-throughput second-generation DNA sequencing (Salas et al., 2017). Our objective was to identify genetic determinants that affected the survival of *E. meliloti* during the industrial production PI formulations. To achieve that objective, sterile peat bags were provided by BIAGRO-BAYER, as well as the facilities for postinoculation storage.

Fig. 1 summarizes the treatments and the experimental approach used. Briefly, each of the 30 mutant sets was grown up to OD_600_ in the range of 2–5, and a dilution (1:8) in peat diluent solution was then made. A part of these cultures was saved as the input sample. Next, 5 ml containing approximately 1.10^9^ CFU/ml were inoculated into sterile peat bags. These bags were taken to BIAGRO-BAYER for the storage period, which was chosen to be of 2 months after taking into account the time required for peat maturation (Materon and Weaver, 1985), the beginning of the rhizobium-population decline, and the period of storage before use. After this time, the surviving bacteria were recovered from the inoculant bags and stored for DNA isolation. A postrecovery growth step in rich medium was required (approximately 4–5 generations) since otherwise the PCR amplification of the DNA samples for Illumina sequencing was not possible. The DNA isolation and qPCR reactions were performed as in Salas et al. (2017) for both the input and the output samples. The PCR products for all the conditions and mutant sets were then mixed in equal amounts, and the final mixture was sequenced with the ILLUMINA HiSeq platform. Since any given Tn5 mutant bore two specific DNA-signature tags (namely, the K and H tags; Pobigaylo et al., 2006), the relative amount of each mutant in any given sample was quantified by the high-throughput DNA sequencing of the bar-coded PCR products individually contained each one of the Tn5 signature tags. A total of 108,379,479 reads were obtained and demultiplexed through the use of the STM-seq-count perl script. The raw counts were separated by pool, input or output sample, K or H signature and biological replicate. For the comparison of the paired input and output samples, the raw counts were normalized by taking the proportions of each mutant in each sample.

**Fig. 1.**
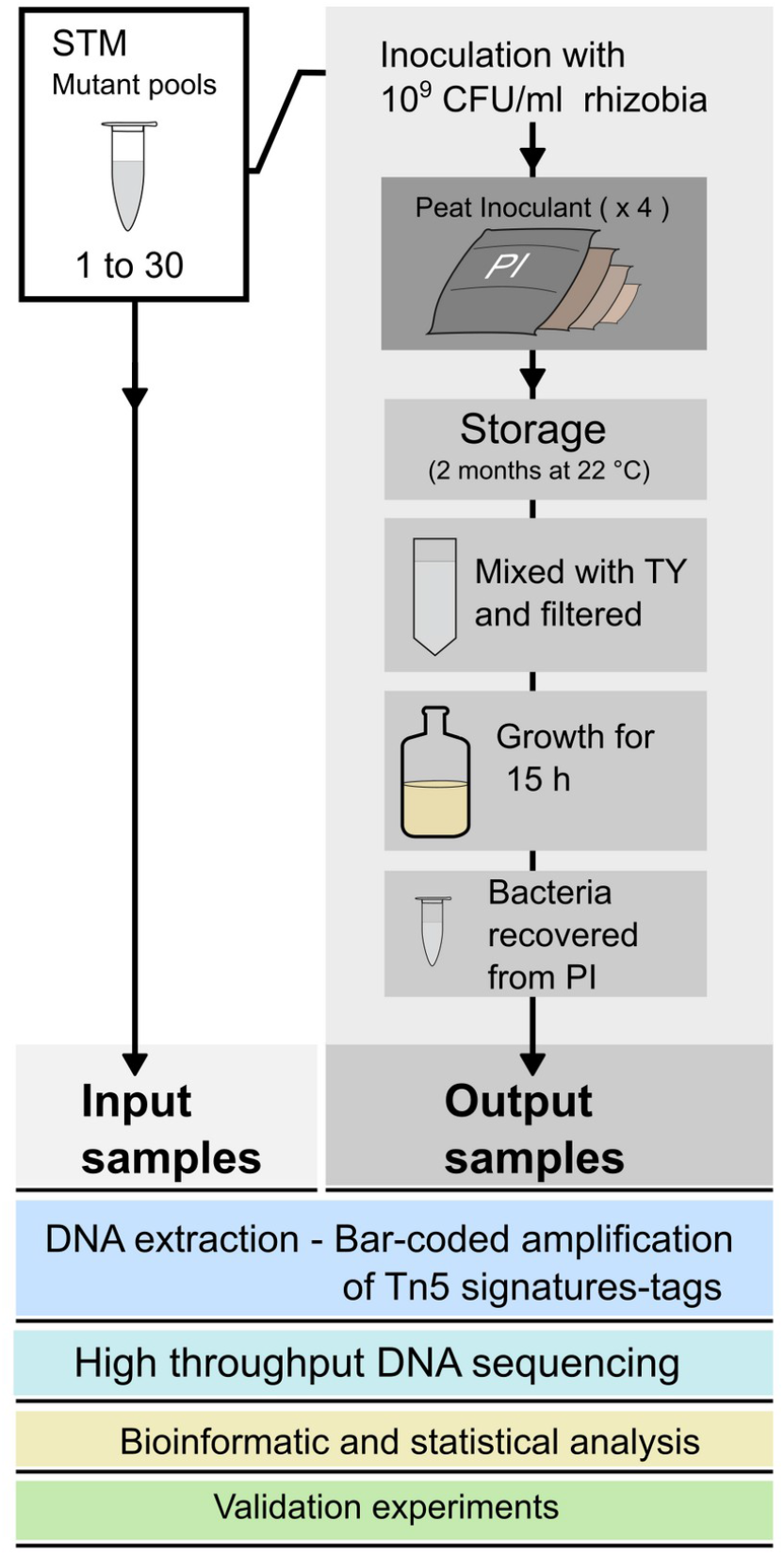
General experimental design for the identification of *Ensifer meliloti* Tn5 mutants affected during the maturation and storage period on peat based inoculant formulations. The scheme summarizes the experimental design used to evaluate the fitness of more than 12,000 Tn5 mutants distributed in 30 sets of *ca*. 412 mutants each (Pobigaylo et al., 2006) during the storage and maturation of peat based inoculant formulations. Plastic bags containing sterile peat were inoculated with *ca*. 10^9^ CFU/mL of rhizobia (input sample). The proportion of each mutant in the input and output samples—the latter recovered two months post inoculation—was estimated by the bar-coded PCR of specific Tn5 signatures followed by high-throughput DNA sequencing (Illumina-HiSeq) of the amplified products (*cf*. details in Materials and Methods). For each set of mutants, under any of the indicated experimental conditions, 4 biological replicates were included. A decreased (or augmented) abundance of a specific mutant in the output sample compared to that of the same mutant in the inoculum was indicative of a Tn5 insertion that possibly affected rhizobial fitness during the treatment.

To assess which mutant rhizobium manifested an altered fitness in PI, the log_2_ of the ratio of the number of each mutant in the output sample with respect to the same mutant in the input sample (inoculum)—*i*.*e*., the M value—was determined by means of homemade scripts (Salas et al., 2017). MA plots were then generated that revealed significant biases in the results, mainly, the average M-value was smaller than 0, and a correlation between the M and A values was observed (Fig. 2.A). These biases are partially the result of the presence of extremely high read counts for a subset of mutants in most of the output samples, and could be explained by the necessity of growing the recovered bacteria prior to the DNA isolation. Of interest to us was that those mutants exhibiting higher read counts either had a mutation that greatly improved their survival on the inoculant, or were less stressed and presented a shorter lag phase, thus outgrowing the rest of the mutants during the postrecovery growth step. In order to correct the observed biases and accurately predict the mutants with an altered fitness phenotype, the IDEGES/EDGER and IDEGES/DESEQ2 pipelines from the TCC R package for differential expression analysis (Sun et al., 2013; *cf*. Materials and Methods) were used. The MA and volcano plots of Fig. 2.B illustrate that the M-value bias was corrected by this approach, though some of the distortion caused by the extreme counts still remained. A mutant was considered to be altered when, under any given condition, both of the signatures (the H and the K tags) displayed a DEG flag in both TCC pipelines (*i*.*e*., IDEGES/EDGER and IDEGES/DESEQ2), an M-value less than −1 (*i*.*e*., at least half the number of mutant counts in the output sample) or greater than 2 (*i*.*e*., four times the number of mutant counts in the output sample), and the sum of the total reads for the mutant (*i*.*e*., input plus output sample reads) was greater than 300. The upper cutoff for the M-value was imposed to be higher than the lower cutoff, because a small bias towards positively affected mutants remained after the normalization (0.2 units on average, Table 2S, Fig. 2.B).

**Fig. 2.**
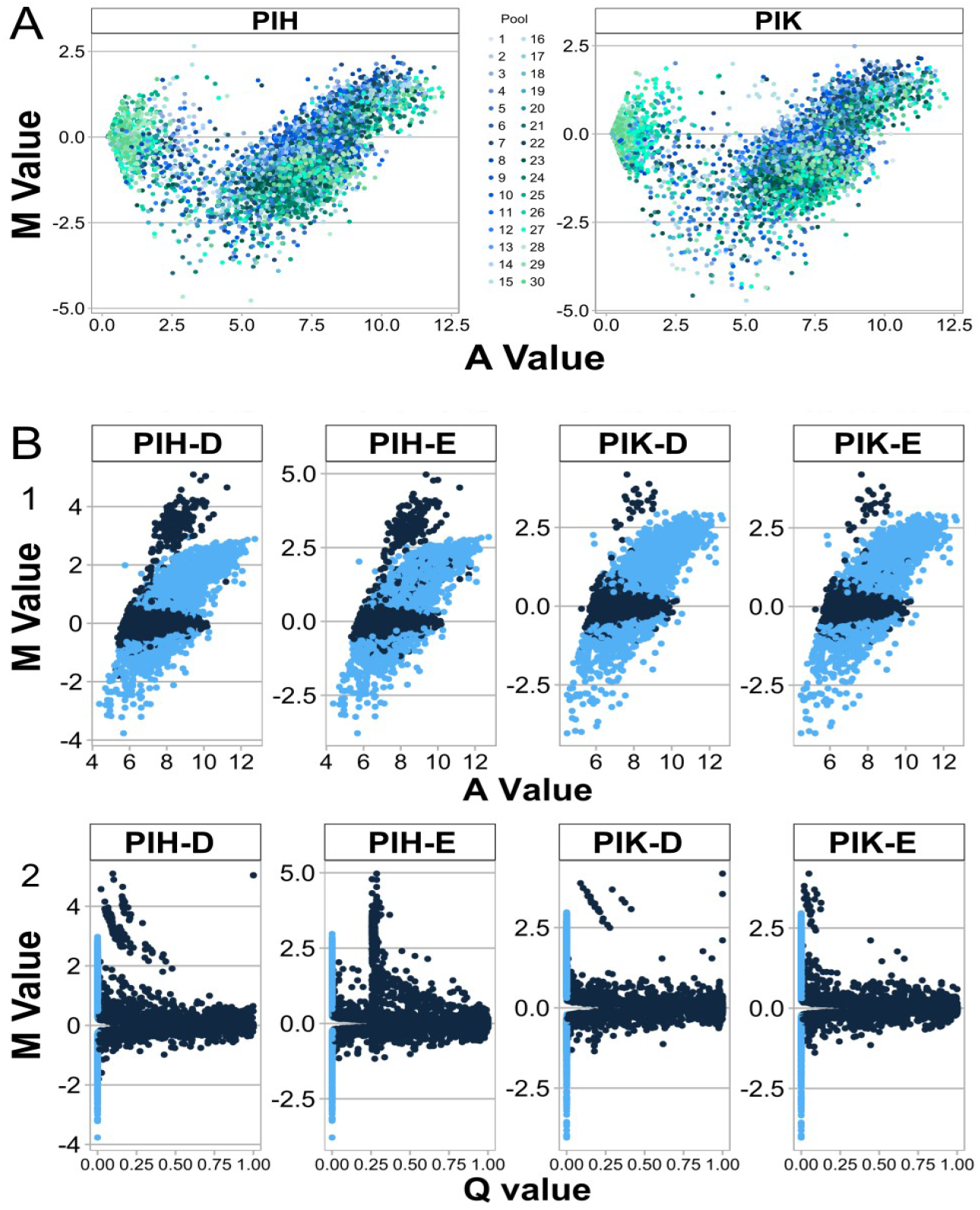
MA and volcano plots for differentially expressed genes during peat inoculant maturation. A) M and A were calculated from the proportions of each mutant under the input and output conditions by means of homemade scripts. Each dot represents a mutant. The colors identify mutants from different STM mutant-pools. B) Normalization of mutant counts and the identification of mutants with an altered phenotype through the use of the TCC R package for differential expression analysis. The M, A and Q values (*i*.*e*., FDR adjusted p-values) calculated by the iDeges/Edger and iDeges/Deseq2 pipelines were used to generate 1) MA and 2) volcano plots. Each dot represents a mutant. The light blue dots represent mutants with predicted differential fitness. The peat inoculant (PI) in combination with H (the H STM signature)—PIH—or K (the K STM signature)—PIK—or with D (the Deseq2 pipeline)—PIH-D and PIK-D —or E (the Edger pipeline)—PIH-E and PIK-E—is indicated, respectively, from left to right above each plot.

On the basis of these criteria, of a total of 12,070 analyzed mutants, 471 (of which 393 have a known insertion site) could be identified as potentially altered during maturation and storage on PI formulations (Table S3). An analysis of the insertion sites of these mutants revealed that more than three hundred different genomic regions were affected, in some instances carrying several insertions within the same gene and/or operon (thus constituting a redundancy situation; Table S2).

### 3.2. Individual phenotypic evaluation of mutants with altered fitness in peat-based inoculants

In order to assess whether mutants identified by STM consistently manifested an alteration in fitness during the PI treatments, the proportion of viable cells after PI treatment was determined for individually selected mutants. To that end, the survival of individual mutants in PI was tested, and although no significant differences were found, most of the mutants with a negative M-value exhibited a smaller relative proportion after the PI treatment than the wild-type strain *E. meliloti* 2011 (Fig. 3).

**Fig. 3.**
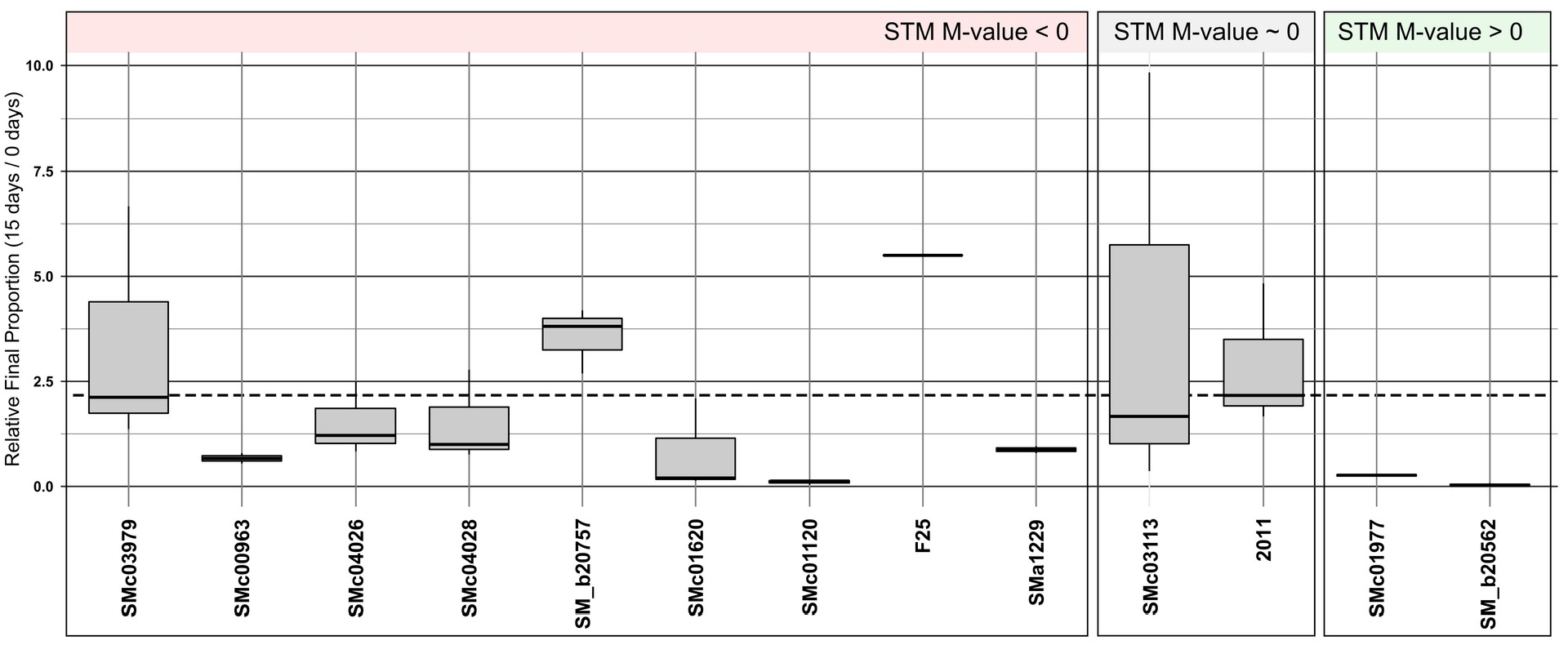
Individual phenotype validation of thirteen selected mutants that manifested altered fitness during survival in PI. The survival in peat inoculant (PI) of thirteen isolated mutants, selected by their M-values (smaller than 1 or greater than 2) in our STM-seq experiment, was evaluated as described in Materials and Methods. The relative final proportion was calculated as the CFUs recovered after 15 days PI treatment with respect to the CFUs recovered at the time of inoculation. Green and light red shadings correspond to mutants with M > 2 or M < −1 respectively. 2011: *Ensifer meliloti* 2011 (Wild type). SMc03113: control, STM mutant presenting near zero M-value. F25: an intergenic Tn5 insertion at position 682,937 in the chromosome. The horizontal broken line represents the relative proportion for the wt strain. In the box plot, the upper and lower borders represent the quartiles of the data, the solid line the median, and the upper and lower outriders (the whiskers) the respective maximum and minimum values.

### 3.3. Genomic and functional distribution of genetic markers involved in the fitness of E. meliloti in PI formulations

The genomic distribution of the Tn5 insertions that affected fitness in PI treatments indicated that nearly 58% were located in the *E. meliloti* chromosome [Fig. 4.B, Mutants (Percent)]. When only the mutations that produced a negative fitness were analyzed, however, the proportion of chromosomal insertions was nearly 84% [Fig. 4.B, Mutants (Percent); PI M < 0]. In the case of mutants that exhibited positive fitness, more than half of the insertions were mapped on the plasmids pSymB and pSymA [Fig. 4.B, Mutants (Percent); PI M>0]. This finding was expected, since the pSym plasmids are more related to symbiotic, adaptative, and accessory functions. The average density of genes involved in PI fitness per replicon length is in the relative order: chromosome > pSymB > pSymA (*cf*. the distribution of orange dots in the light red background in Fig.4.A plus Fig. 4.B, Density, Mutants/Kb) for mutants with negative fitness, and in the order pSymB > pSymA > chromosome for mutants with positive fitness (Fig 4.B, Density, Mutants/Kb). In addition, most of the mutants with an altered fitness phenotype are conserved in the genus *Ensifer (i*.*e*., present in the core genome, obtained from Salas et al., (2017); that is, more than 66% of the total genes. Table S3), and in some of the most related rhizobia (*i*.*e*., more than 34% in *Bradyrhizobium*, and more than 50% in *Rhizobium* and *Mesorhizobium*).

**Fig. 4.**
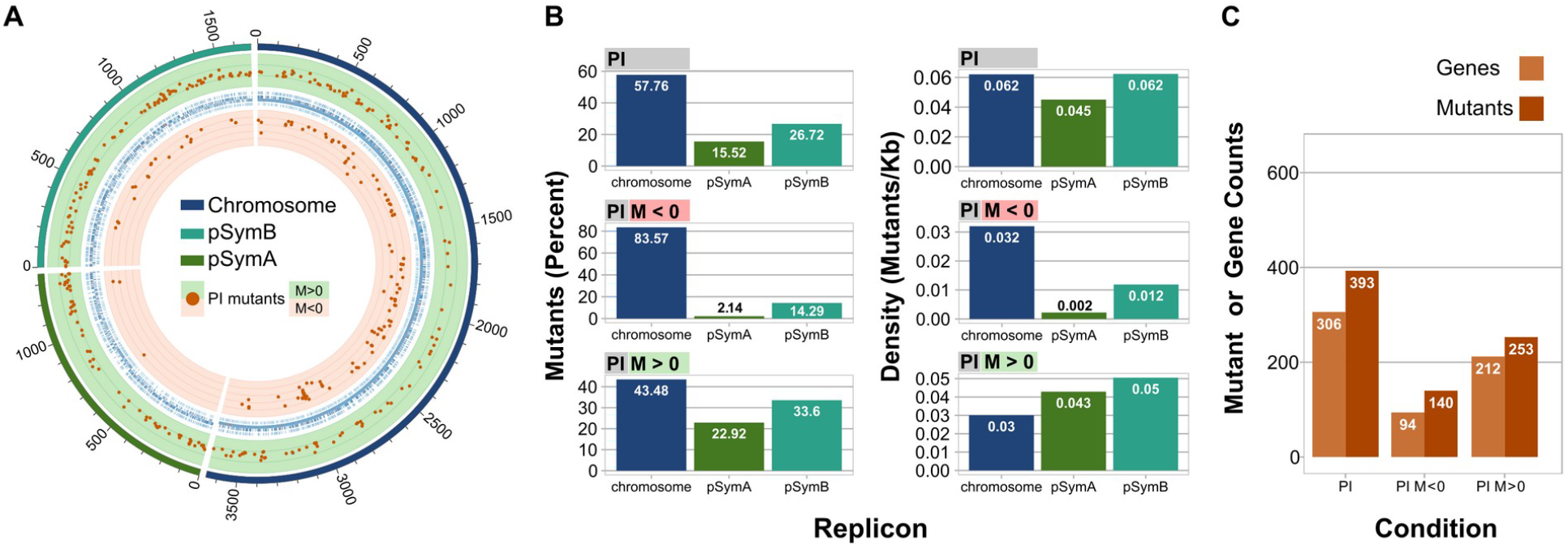
Genomic distribution of *Ensifer meliloti* STM mutants with an altered fitness phenotype on peat based inoculants. A. Circos plot illustrating the position of all the mutants with an altered phenotype. The green and light red backgrounds, correspond to positive fitness or negative fitness, respectively. M: M-value (log2(output count / input count). B. Bar graphs indicating the percent of the mutants located on each replicon (left), or the density of mutants per replicon expressed in mutants per Kb (right). C. Total number of mutants, or genes that after interruption by a transposon, manifested an altered phenotype.

An analysis of the cluster of orthologous genes (COG; Fig. 5, Table S2) indicated that the most affected mutants presented insertions in genes with functions related to the transport and metabolism of amino acids and carbohydrates, transcription, and cellular energy, among others. The COG profiles for the mutations that produced a negative and positive fitness were quite different: in the former, the mutations in the genes related to cell wall and/or membrane biogenesis, amino-acid metabolism and transport, transcription, and energy production and conversion were more relevant; whereas in the latter the most represented categories were carbohydrate metabolism and transport, replication, and recombination and repair. In addition, a high proportion of positive fitness mutants presented insertions in genes with general function prediction, unknown function, or were not found in COGs.

**Fig. 5.**
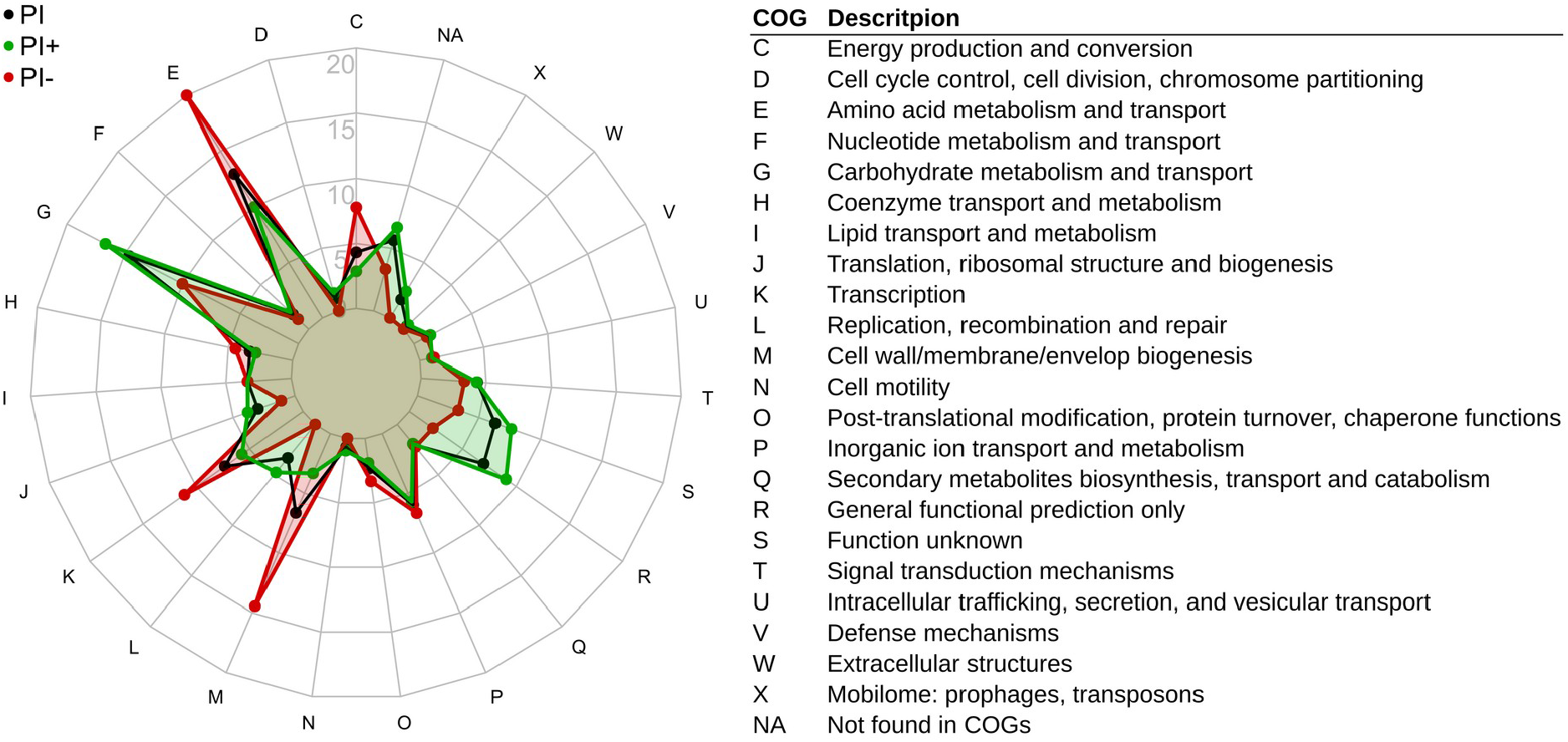
Functional distribution of proteins involved in fitness during PI maturation and storage. Radar diagram of the cluster of orthologous genes (COG) indicating the functional distribution of those proteins in which a mutation generated an altered phenotype on peat based inoculant formulations. PI (Black):all the proteins that when mutated presented an altered fitness phenotype. PI+ (Green): proteins that when mutated generated a fitness advantage. PI- (red): proteins that when mutated generated a fitness disadvantage. The letters in the radar diagram correspond to the COG categories listed to the right of the panel. The radii correspond to the percentage of proteins in each COG category.

### 3.4. Growth response of mutants with altered fitness in PI

To evaluate whether the mutants with differential fitness in the survival during PI maturation were auxotrophs, mutants in 23 selected genes were individually tested for differential growth in rich (TY) and defined (Evans) medium. Growth curves were determined by measuring OD_600_, and the exponential phase growth rates were calculated (Table 1). All the mutants tested grew as the wild type strain in rich medium, but, as expected, certain mutants in genes involved in amino acid (*cysD, ilvC, ilvI, gltD, gltB, metA, pheA*, and *trpF*) and nucleotide (*purL*) biosynthesis were unable to grow in Evans medium. In addition, *zwf*, a part of the pentose phosphate pathway, and *bioN*, a biotin transporter, were also impaired in growth in Evans medium. A finding of interest to us was that certain of the mutants with a fitness advantage on PI, underwent a small increase in the growth rate on Evans medium.

**Table 1.** Exponential-phase growth rates for mutants with an altered PI fitness in rich and defined media.

| Mutant |  | TY | Evans | STM M value (+/-) | Experiment |
| --- | --- | --- | --- | --- | --- |
| 2011 | WT | 0.20 | 0.24 |  | 1 |
| SMb20757 | *bhbA* | 0.19 | 0.23 | - | 1 |
| SMb21463 | - | 0.21 | 0.24 | + | 1 |
| **SMc00488** | *purL* | 0.19 | **0.00** | - | 1 |
| **SMc00641** | *serA* | 0.20 | **0.00** | - | 1 |
| **SMc00963** | *bioN* | 0.18 | **0.10** | - | 1 |
| **SMc01431** | *ilvI* | 0.21 | **0.00** | - | 1 |
| SMc02228 | *fadA* | 0.19 | 0.23 | - | 1 |
| SMc02450 | *argJ* | 0.22 | 0.24 | - | 1 |
| **SMc02767** | *trpF* | 0.25 | **0.04** | - | 1 |
| **SMc02899** | *pheA* | 0.19 | **0.01** | - | 1 |
| **SMc03797** | *metA* | 0.23 | **0.01** | - | 1 |
| **SMc04026** | *gltD* | 0.22 | **0.06** | - | 1 |
| **SMc04028** | *gltB* | 0.20 | **0.06** | - | 1 |
| SMc04331 | *mtbC* | 0.21 | 0.23 | - | 1 |
| **SMc04346** | *ilvC* | 0.26 | **0.01** | - | 1 |
| **SMc04346** | *ilvC* | 0.25 | **0.01** | - | 1 |
| 2011 | WT | 0.27 | 0.21 |  | 2 |
| SMb20757 | *bhbA* | 0.25 | 0.19 | - | 2 |
| 2011 | WT | 0.28 | 0.28 |  | 3 |
| SMb20652 | *asnB* | 0.27 | 0.29 | + | 3 |
| **SMc00091** | *cysD* | 0.27 | **0.03** | - | 3 |
| **SMc01431** | *ilvI* | 0.30 | **0.00** | - | 3 |
| SMc02875 | - | 0.33 | 0.32 | - | 3 |
| **SMc02899** | *pheA* | 0.28 | **0.02** | - | 3 |
| **SMc03070** | *zwf* | 0.27 | **-0.01** | - | 3 |
| **SMc04042** | - | 0.21 | **0.03** | - | 3 |
| 2011 | WT | 0.22 | 0.21 |  | 4 |
| SMc03931 | *soxA2* | 0.22 | 0.20 | - | 4 |
| 2011 | WT | 0.29 | 0.27 |  | 5 |
| SM_b21535 |  | 0.30 | 0.31 | + | 5 |
| SMc04245 | *znuA* | 0.28 | 0.30 | - | 5 |

\*Mutants in bold are those that presented a decrease in the exponential-phase growth rate in Evans medium.

## 4. Discussion

### 4.1. Proteins involved in the survival of E. meliloti in PI maturation and storage

#### 4.1.1. Mutants with decreased fitness

Of the 471 Tn5 insertion mutants that presented an altered phenotype in PI, 176 evidenced a decreased fitness. Most of these mutants contained insertions in genes with an annotated function related to membrane and metabolic processes, mainly amino acid and carbohydrate transport and metabolism (Fig. 5, Table S5).

Of the mutants with negative M-value, the most highly affected bore an insertion in *SMc00828-mutT* (8-oxo-dGTP pyrophosphatase; Table 2). MutT hydrolyses 8-oxo-dGTP to the nucleoside diphosphate, removing the latter from the triphosphate pool in order to prevent its incorporation into the DNA, which would produce a higher frequency A→C transversions (Fowler et al., 1994; Galperin et al., 2006; Maki and Sekiguchi, 1992). *Escherichia coli* MutT mutant*s* have exhibited a greater mutation rate than wild-type strains under both aerobic and anaerobic conditions (Fowler et al., 1994), and strikingly, during starvation (Bridges, 1996), which could explain the important role of MutT during the storage and maturation of inoculant formulations.

**Table 2.**
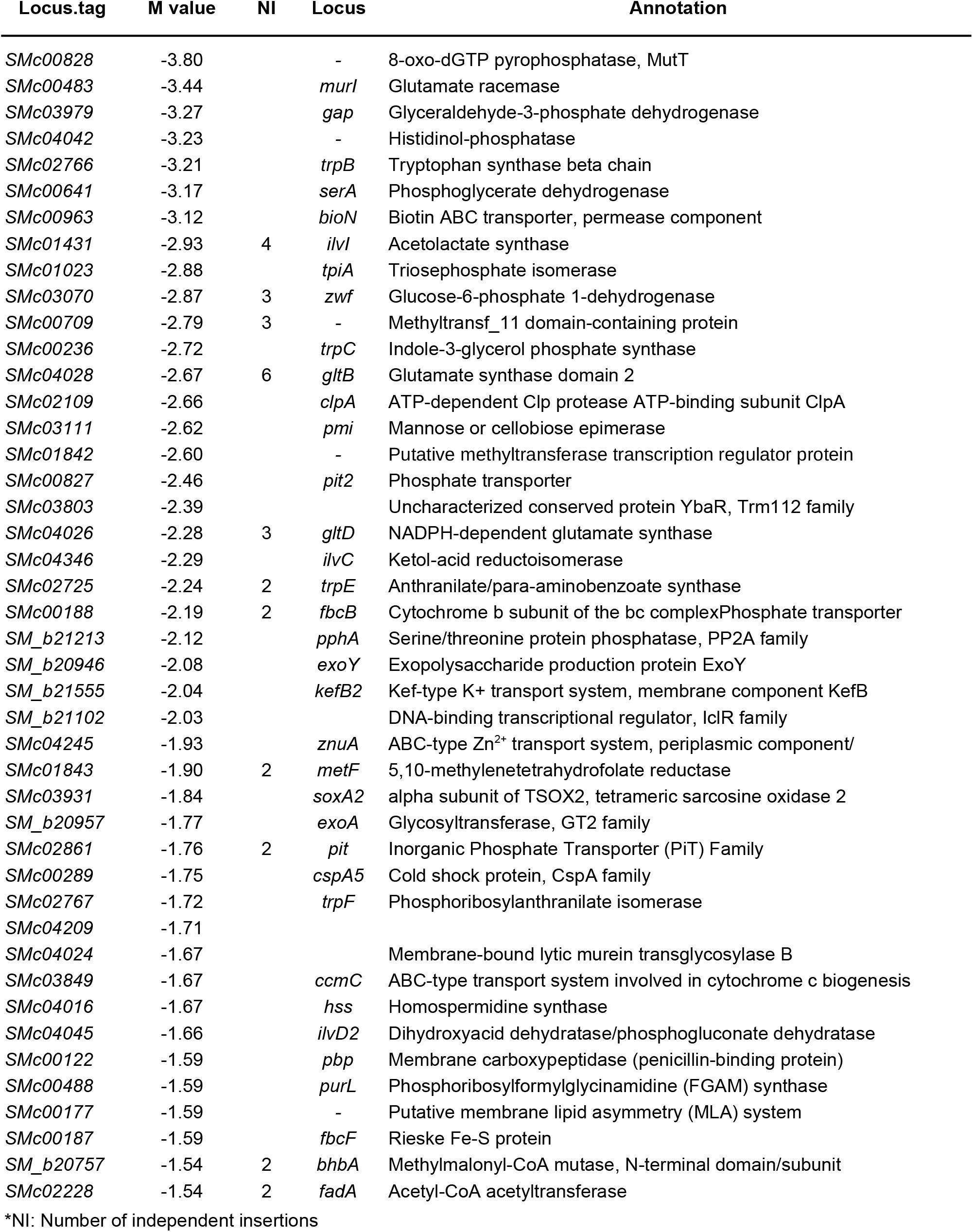
Genes with the most negative M-values (M < −1.5) on PI formulations.

##### Cell envelope

Several mutants with insertions in genes related to cell wall and/or membrane biogenesis were also severely affected. Among those genes, *SMc00483-murI* encodes an enzyme (glutamate racemase) involved in the isomerization of L- and D-glutamate (Doublet et al., 1993), with the latter being a component of the cell-wall peptidoglycan. In addition, MurI from certain bacteria has been found to act as an inhibitor of the DNA gyrase (Sengupta et al., 2008) and has been reported as inducible under acid stress in *E. meliloti* (Draghi et al., 2016). *SMc00122*-*pbp*, another gene affecting rhizobial fitness in PI, encodes for a putative penicillin-binding protein similar to the *E. coli* PBP-1B. These proteins are bifunctional with transglycosylase (at the N terminus) and transpeptidase (at the C terminus) activities, involved in the final stages of the synthesis of peptidoglycan. In a recent publication by Atieno et al. (2018), an aqueous peat extract was used as a model to study peat protein production in *R. leguminosarum* bv viciae and trifolii, and *Bradyrhizobium diazoefficiens*. In that work, a penicillin-binding protein was found to be induced in the rhizobia after the treatment of bacteria with the peat extract. Our results have demonstrated that *SMc04024*, another protein involved in cell wall maintenance and biosynthesis (a putative membrane-bound lytic murein transglycosylase B) was also important for rhizobial survival in PI. All together, the results summarized here underscore the relevance of an adequate peptidoglycan structure and dynamics. With respect to other components of the bacterial envelope, mutants with transposon insertions in the *exoY* and *exoA* genes—*i*.*a*., normally completely devoid of succinoglycan (EPS-I, Davies and Walker, 2007; Long et al., 1988)*—*were also negatively affected in their peat survival. Whereas high osmotic pressure stimulates the synthesis of the high-molecular-weight form of this polymer (Janczarek, 2015; Miller-Williams et al., 2006), an acidic pH produces an increment in the low-molecular-weight EPS-I (Hawkins et al., 2017; Hellweg et al., 2009) and the induction of several genes involved in the EPS-I biosynthesis (Draghi et al., 2016; Hellweg et al., 2009). Nonetheless, still-unconnected evidence for the role of this polysaccharide in the tolerance of the rhizobia to abiotic stresses (Cytryn et al., 2007; Davies and Walker, 2007; Dilworth et al., 1999; Miller-Williams et al., 2006), and possibly to PI, exists. In agreement with previous reports (Berninger et al., 2018), our results indicate that exopolysaccharides (EPS) play a major protective role in inoculant formulations. In addition to EPS-I, other polysaccharides appear to be of relevance for the survival of *E. meliloti* in PI (namely, the capsular polysaccharide, KPS and lipopolysaccharide, LPS) since we have seen that mutants on *rkpZ2, rkpZ1, rkpJ, lpsE*, and lpcC presented a mild fitness decrease. Furthermore, a mutation in *SMc00177*, an ABC-type transport auxiliary lipoprotein component homologous to the *E. coli* membrane lipid asymmetry (MLA) system, was severely affected on PI. This system helps to maintain the lipid asymmetry in the outer membrane of the Gram-negative cell envelope. Arnold et al. (2017), have reported that mutations in the SMc00177 ABC system increase bacterial fitness during the treatment with the nodule-specific-cysteine-rich peptide NCR247, possibly by altering the outer membrane lipid composition. These results suggest that the integrity of the membrane is important for *E. meliloti* survival in PI formulations. Our findings are reinforced by previous reports for *Rhizobium leguminosarum* (Atieno et al., 2018), for which growth in peat extract has been found to expose rhizobia to a sublethal stress, cause damage to cell membranes, and induce the expression of proteins involved in cell membrane maintenance and repair, among others.

##### Amino acid metabolism

Several enzymes involved in amino acid biosynthesis were required for survival on PI, suggesting that during the inoculant maturation an amino acid shortage might be occurring. The most affected mutants contained insertions in *SMc04042*, a gene that was recently described as encoding the missing *E. meliloti* histidinol phosphate phosphatase (diCenzo et al., 2018) required for histidine biosynthesis; *SMc01431-ilvI, SMc04346-ilvC and SMc04045-ilvD2*, required for the biosynthesis of valine, leucine and isoleucine; *SMc00091*-*cysD*, r*e*quired for cysteine biosynthesis; *SMc00641-serA*, involved in serine and cysteine biosynthesis; *SMc02766*-*trpB, SMc00236-trpC, SMc02725-trpE* and *SMc02767-trpF*, r*e*quired for tryptophan biosynthesis; *SMc03797*-*metA and SMc01843-metF*, required for methionine biosynthesis; SMc02717-*leuA1* r*e*quired for leucine biosynthesis; *SMc02899*-*pheA* r*e*quired for phenylalanine biosynthesis; and *SMc02450-argJ*, required for arginine biosynthesis. Moreover, *SMc04028-gltB* and *SMc04026-gltD* involved in nitrogen assimilation were also important for PI fitness. Most of these mutants were unable to grow in a defined minimal medium (Table 1) suggesting that several phenotypes in PI may be due to auxotrophy. Since in *E. meliloti* a betaine-homocysteine methyltransferase was reported to replace MetF as a methyl donor in the presence of glycine-betaine, the observed impairment in the present *metF* mutant suggests that PI likely contains no significant amounts of glycine-betaine (or choline).

##### Lipid and carbohydrate metabolism. Energy production and conversion

Mutants on several proteins related to lipid metabolism proved to be impaired on PI fitness. Those mutants comprised *SMc01620-eryB* (encoding Glycerol-3-phosphate dehydrogenase), which serves as a major link between carbohydrate metabolism and lipid metabolism; *SM_b20757-bhbA* (encoding Methylmalonyl-CoA mutase) required for the degradation of odd-chain fatty acids (and also of valine, isoleucine, methionine, threonine and thymine); and the three genes *SMc02227-fadB, SMc02228-fadA, and SMc02229* (putative acetyl-CoA acyltransferase, putative fatty oxidation complex alpha, and putative acyl-CoA dehydrogenase, respectively) which are involved in lipid anabolism and catabolism. Although *fadA* was reported to be induced under different stress conditions (Krol and Becker, 2011), *fadD—*which was implicated in the long term survival of *E. meliloti* and *E. coli* (Pech-Canul et al., 2011)—quite unexpectedly did not affect rhizobial fitness in PI. Survival in peat required genes related to central carbohydrate metabolism including *SMc03979-gap, SMc01023-tpiA*, and *SMc03111-pmi*, along with others like *SMc02322-rhaD* (a short-chain dehydrogenase/reductase that is part of the rhamnose utilization operon) and *zwf* (a key glucose-6-phosphate-1-dehydrogenase of the pentose phosphate pathway), which in our laboratory was found to be induced under acid stress in *E. meliloti* (Draghi et al., 2016). The inactivation of *zwf* also resulted in an osmosensitive phenotype with loss of osmoprotection by trehalose and sucrose (Barra et al., 2003). The relevance of energy production and conversion to PI survival was supported by the strongly impaired phenotype observed in mutants like *SMc00187-fbcF* and *SMc00188-fbcB*, involved in electron transport and the proton movement across the cell membrane, respectively. Several lines of evidence suggest that FbcB aids in cells survival under stress conditions by providing an auxiliary resource for mobilizing PPi energy and maintaining cation gradients (Baltscheffsky et al., 1999).

##### Transcription regulation and signal transduction mechanisms

Genes in this group included *SM_b21213-pphA*, a gene coding for a putative serine/threonine protein phosphatase (STPs) homologous to PssZ from *Rhizobium leguminosarum* bv trifolii Rt24.2; whose mutation causes pleiotropic effects including the lack of EPS production, reduced growth kinetics and motility, and failure in host root infection (Lipa et al., 2019, 2018). Another key gene for PI survival was *fixL*, the histidine kinase of the FixL-FixJ two-component system. FixL senses oxygen, and when oxygen concentration is low, the phosphate group in FixL can be transferred to the response regulator FixJ, which in the phosphorylated state (FixJ-P), is responsible for the activation of the master regulators of nitrogen fixation, *nifA* and *fixK*. NifA induces the expression of the structural genes encoding the nitrogenase, while FixK induces the expression of a high-affinity terminal oxidase (*cbb*_*3*_ *cytochrome oxidase*) required for bacterial respiration under microaerobic conditions (Rutten and Poole, 2019). That FixL is required for PI survival further supports the notion that a decrease in the oxygen concentration within the PI bags is taking place during the storage and maturation process. Several other transcriptional regulators were found to be associated with a better survival of rhizobia in PI; including *SMc01507-rsiC* which encodes the histidine kinase of a two-component system, plus eight transcriptional regulators (*SMc03140, SMa2287, SMc01842, SMc01164-iolR, SM_b20717, SM_b21299, SM_b21102, SMc02172*-*frcR* and *SM_b20323*) some of which had already been reported to be important for diverse cellular processes including the conversion of exogenous deoxyribonucleosides (Taboada et al., 2012), sugar transport (Lambert et al., 2001), sulfite respiration (Lang et al., 2018), and rhizosphere colonization (Salas et al., 2017).

The rest of the mutants with a negative fitness belong to diverse COG functional categories including genes for purine metabolism (SMc00488-purL), biotin transport (*SMc00963-bioN*, Guillen-Navarro et al., 2005), and phosphate metabolism (*SMc00618-ppk, SMc00827-pit2, SMc02861-pit*). Of interest to us was that, *SMc02861*-*pit*, which encodes a putative Pi transport system that is expressed under excess Pi and repressed under Pi limitation (Bardin et al., 1998), was severely affected in its PI survival. This phenotype suggests that Pi is abundant in peat, and that Pit might be an significant Pi importer under such conditions. Moreover, metabolites that promote osmotolerance also very likely had a positive effect on the rhizobial survival in peat. *SMc03931* is involved in the catabolism of choline, a precursor of glycine-betaine that is a potent osmoprotectant in bacteria (Fougere and Le Rudulier, 1990). The relevance of other genes like *SMc04016-hss* whose product is a probable homospermidine synthase also suggest that rhizobia have to cope with challenging stress conditions in the PI medium.

#### 4.1.2. Mutants with increased fitness

In the case of proteins whose mutation led to a better fitness during the storage and maturation of PI, the more prevalent COG classes were carbohydrate transport and metabolism, amino acid transport and metabolism, and unknown function (Figure 5, Table S5). The positive fitness associated with mutations in genes of certain transporters encoded in the symbiotic megaplasmids suggests a negative cost-to-benefit balance in the expression of these genes in the rhizobia persisting in peat. In the case of amino acid transport, several mutants harbored insertions in genes annotated as permease components of ABC-type transport systems. Likewise, several mutants in carbohydrate transport also included Tn5 insertions in ABC-type transport systems for lactose (*SM_b20969, SM_b20971, SM_b21594*), fucose (*SMa2311*), rhamnose (*SMc02324*), arabinose (*SM_b20895*-*araA*), galactose (*SM_b21343*), ribose (*SM_b21343*), and trehalose-maltose (*SMb20327*-*thuG*, Jensen et al., 2002). With respect to carbohydrate metabolism, the finding that *glgB2* encoding the 1,4-alpha-glucan branching enzyme and *glgP* encoding for glucan phosphorylase are not required for PI fitness suggests that the use of glycogen as an energy source might generate a disadvantage in the long term. Mutants in *aceA* (*SMc00768*) were found with high M values in PI. Since AceA encodes for an isocitrate lyase, the enzyme catalyzing the first step of the glyoxylate cycle, a gluconeogenic carbon source is very likely available during the maturation process. Mutations in genes associated with polysaccharide biosynthesis (including EPS and arabinose-contaning polysaccharides) prompt us to pay attention to the participation of these components in rhizobial survival in PI, but further studies will be necessary to understand if an increased survival is associated with increased or decreased amounts of those components.

Certain mutations in transcriptional regulators and genes involved in signal transduction mechanisms were found, including an extracytoplasmic sigma factor (*SMc04051-rpoE4*), a putative response regulator (*SMb21561*) homologous to CpxR, a well characterized system to sense misfolded proteins (Santos et al., 2010); a functional diguanylate cyclase with a GGDEF motif (*SMc01464;* Santos et al., 2010); an adenylate cyclase (*SMc02176*-*cyaD1*) induced within nodules (Tian et al., 2012; Zou et al., 2017); and several transcriptional regulators of unknown function.

Other mutants with positive M values contained Tn5 insertions in genes associated with recombination and repair (*SMa0417, SMc01125, SMa2257, SMc03877*); translation related functions (*SMc00826, SM_b20024, SMc00973, SMc02799*); and inorganic ion transport and metabolism, particularly iron [*SM_b20363, SMc00371, SMa1740, SMc02889*, and *SMa2406* genes encoding a Fe^3+^ ABC transporter, a ferritin-like protein, a siderophore-interacting protein, a putative ferrisiderophore permease (Cuív et al., 2004; Viguier et al., 2005), and an enzyme involved in rhizobactin siderophore biosynthesis (Lynch et al., 2001), respectively], copper (*SM_b21578*), protons (*SMa0185*), potasium (*SMa2331*), and nitrate (*SMa0583*). Moreover, mutants in genes related to nucleotide metabolism (*SM_b20128, SMa0101, SMc01821*), along with genes associated with secondary metabolism (*SMa0034*, Kida et al., 2007; *SM_b21640*) also manifested a positive fitness. A mutant on *mcpX* (*SMc01104*), the chemoreceptor involved in chemotaxis towards quaternary ammonium compounds also exhibited positive fitness, suggesting that those compounds are probably not necessary in the long term rhizobial persistence in PI. Finally, many mutants harbored insertions in genes belonging to less represented COGs; or within the general function prediction, and the unknown function categories.

## 5. Conclusions

In this work we used STM-seq to search for genes with a relevant role in the rhizobial survival during the maturation and storage of peat-based inoculants. The functional profile of the identified genes suggests that peat is a stressing sublethal environment for rhizobia, with disturbing factors that likely include suboptimal osmotic and nutritional conditions, and oxidative stress. Several genes known to be involved in the stress response, or in starvation as well as in the stationary growth phase, were required for rhizobial survival in PI. In agreement with previous observations indicating that exposure of rhizobia to peat extracts produced membrane damage (Atieno et al., 2018), several cell wall biosynthetic and/or modifying enzymes were found to be required for a better survival of *E. meliloti* in PI. Certain of the markers identified serve to make some inferences related to specific compounds that could be evaluated as additives for the improvement of PI formulations.

Furthermore, these results enabled us to make inferences about the availability of nutrients in the PI that could be used as a basis for the improvement of PI formulations. A strong observation was that most of the enzymes involved in amino acid biosynthesis were required, while many transporters involved in amino acid uptake were not—and were even associated with positive M-values likely due to an unnecessary biosynthetic cost of those proteins—thus clearly indicating that free amino acids were not available in PI. That MetF was required could indicate that glycine-betaine (or its precursor choline) is either unavailable. or present in insufficiently high amounts for betaine-homocysteine methyltransferase (*bmt*) to functionally replace MetF in the methionine biosynthesis. In addition, the negative fitness of mutants in the *rhaD* and *thuA* genes, within the rhamnose and trehalose catabolism operons, suggests that when the sucrose added to the PI is consumed, the ability to use those sugars might become an advantage. In contrast, mutants in several transcriptional regulators, in particular the putative repressors of operons involved in the catabolism of nutrients that might not be available in PI, represented a significant disadvantage. The requirement of several enzymes linked to the fatty acid catabolism could indicate that lipids provide an important source of energy in PI. With respect to inorganic ions, ZnuA was required for an appropriate fitness, suggesting that PI contains a low concentration of Zn^2+^. This conclusion was further supported by the positive fitness presented by *SMc04128* mutants, which are required for tolerance to high concentrations of Zn. That mutants in the inorganic-phosphate transporter PiT—which is expressed only under high phosphate concentrations and repressed under phosphate limitation—manifested a decreased fitness suggests that inorganic phosphate is abundant on PI. Of particular relevance is that several mutants in genes involved in the uptake of Fe, Mg^2+^, Cu^+^, K^+^, and NO_3-_ presented positive M-values, indicating that PI likely contains sufficient amounts of all these ions with a bioavailability that is sufficient to satisfy the requirement of the inoculated rhizobia.

As to rhizobial respiration, the requirement of *fbcF*, which is a part of the cytochrome bc1 complex (complex III) induced under microaerobic conditions, and *fixL*, the membrane-bound oxygen sensor, both suggest a low oxygen concentration during the long time periods used for PI maturation. Finally, a mutant in MutT, a protein required to prevent the incorporation of 8-oxo-dGTP, which generates an increased rate of transversions in DNA, was the most greatly affected in PI. *Echerichia coli* MutT mutants present a higher mutation rate during starvation (Bridges, 1996) a condition that may be operative during the PI maturation. Under such conditions, an active MutT likely has a relevant role in preventing the generation of lethal mutations.

The functional diversity associated with genes affecting the long-term survival of rhizobia in PI constitutes evidence in support of the notion that persistence in that medium involves a complex phenotype that is connected to diverse cellular activities, mostly related to satisfying the requirements of bacterial nutrition (*e*.*g*., *carbon* sources, ions) and to coping with specific stresses (*e*.*g*., oxidative, mutational). The phenomic approach that we used in this work proved to be a powerful tool for identifying the list of rhizobial genes required for the bacterial survival during PI maturation, together with an index that served to rank the various degrees of relevance of those genes.

The markers identified here constitute an initial but substantial input into the design and implementation of practical strategies based on both the potential addition of components with positive effects for the survival of rhizobia, and the characterization of better preservation conditions that would be compatible with the responses that we now know to occur in the rhizobia during the PI maturation.

## CrediT authorship contribution statement

M. J. Lozano: Conceptualization, Methodology, Validation, Investigation, Bioinformatic analysis, Writing (original draft and review & editing). Ezequiel, G. Mogro: Investigation, Validation, Writing (review & editing). M. Eugenia Salas: Conceptualization, Methodology, Investigation, Validation, Writing (original draft and review & editing). Sofía A. Erdozain: Investigation, Writing(review). Nicolás E. Zuber: Investigation, Writing(review). Anke Becker: Methodology, Writing (review & editing), Funding acquisition. A. Lagares: Conceptualization, Methodology, Investigation, Writing (original draft and review & editing), Funding acquisition.

## Acknowledgments

This research was supported by PICT-2017-2022, and PICT-2019-1021 from MinCyT (Ministry of Science Technology and Productive Innovation, Argentina), PUE from CONICET (National Scientific and Technical Research Council, Argentina), DAAD (Deutscher Akademischer Austauschdienst, Germany) and AvH (Alexander von Humboldt Foundation, Germany).

M.J.L., E.G.M, M.E.S, and N.E.Z were supported by CONICET. S.A.E. was supported by Agencia-MinCyT, and A.L. was supported by both CONCET and the UNLP (Universidad Nacional de La Plata, Argentina).

We are grateful to Paula Giménez, Silvana Tongiani, Juan Guzmán Cook (all members of CPA CONICET at IBBM), and Ruben Bustos (from UNLP) for their technical assistance. Donald F. Haggerty edited the final version of the manuscript.

## References

Albareda, M., Rodríguez-Navarro, D.N., Camacho, M., Temprano, F.J., 2008. Alternatives to peat as a carrier for rhizobia inoculants: Solid and liquid formulations. Soil Biol. Biochem. 40, 2771–2779. https://doi.org/10.1016/j.soilbio.2008.07.021

Andrews, S., 2010. FastQC: A quality control tool for high throughput sequence data.

Arnold, M.F.F., Shabab, M., Penterman, J., Boehme, K.L., Griffitts, J.S., Walker, G.C., 2017. Genome-Wide Sensitivity Analysis of the Microsymbiont Sinorhizobium meliloti to Symbiotically Important, Defensin-Like Host Peptides. MBio 8, e01060–17. https://doi.org/10.1128/mBio.01060-17

Atieno, M., Wilson, N., Casteriano, A., Crossett, B., Lesueur, D., Deaker, R., 2018. Aqueous peat extract exposes rhizobia to sub-lethal stress which may prime cells for improved desiccation tolerance. Appl. Microbiol. Biotechnol. 102, 7521–7539. https://doi.org/10.1007/s00253-018-9086-2

Baltscheffsky, M., Schultz, A., Baltscheffsky, H., 1999. H+-PPases: A tightly membrane-bound family. FEBS Lett. https://doi.org/10.1016/S0014-5793(99)90617-8

Bardin, S.D., Voegele, R.T., Finan, T.M., 1998. Phosphate Assimilation in Rhizobium (Sinorhizobium) meliloti: Identification of a pit-like gene. J. Bacteriol. 180, 4219–4226. https://doi.org/10.1128/jb.180.16.4219-4226.1998

Barra, L., Pica, N., Gouffi, K., Walker, G.C., Blanco, C., Trautwetter, A., 2003. Glucose 6-phosphate dehydrogenase is required for sucrose and trehalose to be efficient osmoprotectants in Sinorhizobium meliloti. FEMS Microbiol. Lett. 229, 183–188.

Beringer, J.E., 1974. R Factor Transfer in Rhizobium leguminosarum. J. Gen. Microbiol. 84, 188–198. https://doi.org/10.1099/00221287-84-1-188

Berninger, T., González López, Ó., Bejarano, A., Preininger, C., Sessitsch, A., 2018. Maintenance and assessment of cell viability in formulation of non-sporulating bacterial inoculants. Microb. Biotechnol. 11, 277–301. https://doi.org/10.1111/1751-7915.12880

Bishop, A.H., Rachwal, P.A., 2014. Identification of genes required for soil survival in Burkholderia thailandensis by transposon-directed insertion site sequencing. Curr. Microbiol. 68, 693–701. https://doi.org/10.1007/s00284-014-0526-7

Bishop, A.H., Rachwal, P.A., Vaid, A., 2014. Identification of genes required by Bacillus thuringiensis for survival in soil by transposon-directed insertion site sequencing. Curr. Microbiol. 68, 477–485. https://doi.org/10.1007/s00284-013-0502-7

Blankenberg, D., Kuster, G. Von, Coraor, N., Ananda, G., Lazarus, R., Mangan, M., Nekrutenko, A., Taylor, J., 2010. Galaxy: A web-based genome analysis tool for experimentalists. Curr. Protoc. Mol. Biol. Chapter 19, Unit 19.10.1-21. https://doi.org/10.1002/0471142727.mb1910s89

Bridges, B.A., 1996. Elevated mutation rate in mutT bacteria during starvation: Evidence for DNA turnover? J. Bacteriol. 178, 2709–2711. https://doi.org/10.1128/jb.178.9.2709-2711.1996

Brockwell, J., Bottomley, P.J., 1995. Recent advances in inoculant technology and prospects for the future. Soil Biol. Biochem. 27, 683–697. https://doi.org/10.1016/0038-0717(95)98649-9

Casteriano, A., Wilkes, M.A., Deaker, R., 2013. Physiological Changes in Rhizobia after Growth in Peat Extract May Be Related to Improved Desiccation Tolerance. Appl. Environ. Microbiol. 79, 3998–4007. https://doi.org/10.1128/AEM.00082-13

Catroux, G., Hartmann, A., Revellin, C., 2001. Trends in rhizobial inoculant production and use. Plant Soil 230, 21–30. https://doi.org/10.1023/A:1004777115628

Cuív, P.Ó., Clarke, P., Lynch, D., O’Connell, M., 2004. Identification of rhtX and fptX, Novel Genes Encoding Proteins That Show Homology and Function in the Utilization of the Siderophores Rhizobactin 1021 by Sinorhizobium meliloti and Pyochelin by Pseudomonas aeruginosa, Respectively. J. Bacteriol. 186, 2996–3005. https://doi.org/10.1128/JB.186.10.2996-3005.2004

Cytryn, E.J., Sangurdekar, D.P., Streeter, J.G., Franck, W.L., Chang, W.S., Stacey, G., Emerich, D.W., Joshi, T., Xu, D., Sadowsky, M.J., 2007. Transcriptional and physiological responses of Bradyrhizobium japonicum to desiccation-induced stress. J. Bacteriol. 189, 6751–6762. https://doi.org/10.1128/JB.00533-07

Dart, P.J., Roughley, R.J., Chandler, M.R., 1969. Peat Culture of Rhizobium trifolii : an Examination by Electron Microscopy. J. Appl. Bacteriol. 32, 352–357. https://doi.org/10.1111/j.1365-2672.1969.tb00983.x

Davies, B.W., Walker, G.C., 2007. Identification of Novel Sinorhizobium meliloti Mutants Compromised for Oxidative Stress Protection and Symbiosis. J. Bacteriol. 189, 2110–2113. https://doi.org/10.1128/JB.01802-06

de Moraes, M.H., Desai, P., Porwollik, S., Canals, R., Perez, D.R., Chu, W., McClelland, M., Teplitski, M., 2017. Salmonella persistence in tomatoes requires a distinct set of metabolic functions identified by transposon insertion sequencing. Appl. Environ. Microbiol. 83, 1–18. https://doi.org/10.1128/AEM.03028-16

Deaker, R., Roughley, R.J., Kennedy, I.R., 2004. Legume seed inoculation technology - A review. Soil Biol. Biochem. 36, 1275–1288. https://doi.org/10.1016/j.soilbio.2004.04.009

diCenzo, G.C., Benedict, A.B., Fondi, M., Walker, G.C., Finan, T.M., Mengoni, A., Griffitts, J.S., 2018. Robustness encoded across essential and accessory replicons of the ecologically versatile bacterium Sinorhizobium meliloti. PLoS Genet. 14, e1007357. https://doi.org/10.1371/journal.pgen.1007357

Dilworth, M.J., Rynne, F.G., Castelli, J.M., Vivas-Marfisi, A.I., Glenn, A.R., 1999. Survival and exopolysaccharide production in Sinorhizobium meliloti WSM419 are affected by calcium and low pH. Microbiology 145, 1585–1593. https://doi.org/10.1099/13500872-145-7-1585

Doublet, P., Van Heijenoort, J., Bohin, J.P., Mengin-Lecreulx, D., 1993. The murI gene of Escherichia coli is an essential gene that encodes a glutamate racemase activity. J. Bacteriol. 175, 2970–2979. https://doi.org/10.1128/jb.175.10.2970-2979.1993

Draghi, W.O., Del Papa, M.F., Hellweg, C., Watt, S.A., Watt, T.F., Barsch, A., Lozano, M.J., Lagares, A., Salas, M.E., López, J.L., Albicoro, F.J., Nilsson, J.F., Torres Tejerizo, G.A., Luna, M.F., Pistorio, M., Boiardi, J.L., Pühler, A., Weidner, S., Niehaus, K., Lagares, A., 2016. A consolidated analysis of the physiologic and molecular responses induced under acid stress in the legume-symbiont model-soil bacterium Sinorhizobium meliloti. Sci. Rep. 6, 29278. https://doi.org/10.1038/srep29278

Duong, D.A., Jensen, R. V., Stevens, A.M., 2018. Discovery of Pantoea stewartii ssp. stewartii genes important for survival in corn xylem through a Tn-Seq analysis. Mol. Plant Pathol. 19, 1929–1941. https://doi.org/10.1111/mpp.12669

Evans, C.G.T., Herbert, D., Tempest, D.W., 1970. Chapter XIII The Continuous Cultivation of Micro-organisms: 2. Construction of a Chemostat. Methods Microbiol. 2, 277–327. https://doi.org/10.1016/S0580-9517(08)70227-7

Feng, L., Roughley, R.J., Copeland, L., 2002. Morphological changes of rhizobia in peat cultures. Appl. Environ. Microbiol. 68, 1064–70. https://doi.org/10.1128/aem.68.3.1064-1070.2002

Fougere, F., Le Rudulier, D., 1990. Glycine betaine biosynthesis and catabolism in bacteroids of Rhizobium meliloti: effect of salt stress. J. Gen. Microbiol. 136, 2503–2510. https://doi.org/10.1099/00221287-136-12-2503

Fowler, R.G., Erickson, J.A., Isbell, R.J., 1994. Activity of the Escherichia coli mutT mutator allele in an anaerobic environment. J. Bacteriol. 176, 7727–7729. https://doi.org/10.1128/jb.176.24.7727-7729.1994

Galperin, M.Y., Moroz, O. V., Wilson, K.S., Murzin, A.G., 2006. House cleaning, a part of good housekeeping. Mol. Microbiol. 59, 5–19. https://doi.org/10.1111/j.1365-2958.2005.04950.x

Goodman, A.L., McNulty, N.P., Zhao, Y., Leip, D., Mitra, R.D., Lozupone, C.A., Knight, R., Gordon, J.I., 2009. Identifying Genetic Determinants Needed to Establish a Human Gut Symbiont in Its Habitat. Cell Host Microbe 6, 279–289. https://doi.org/10.1016/j.chom.2009.08.003

Gourion, B., Berrabah, F., Ratet, P., Stacey, G., 2014. Rhizobium–legume symbioses: the crucial role of plant immunity. Trends Plant Sci. 20, 186–194. https://doi.org/10.1016/j.tplants.2014.11.008

Graham, P.H., Vance, C.P., 2000. Nitrogen fixation in perspective: an overview of research and extension needs. F. Crop. Res. 65, 93–106. https://doi.org/10.1016/s0378-4290(99)00080-5

Guillen-Navarro, K., Araiza, G., Garcia-de los Santos, A., Mora, Y., Dunn, M.F., Guille, K., Mora, Y., Dunn, M.F., 2005. The Rhizobium etli bioMNY operon is involved in biotin transport. FEMS Microbiol Lett 250, 209–219. https://doi.org/S0378-1097(05)00481-7 [pii]10.1016/j.femsle.2005.07.020

Hawkins, J.P., Geddes, B.A., Oresnik, I.J., 2017. Succinoglycan production contributes to acidic ph tolerance in Sinorhizobium meliloti Rm1021. Mol. Plant-Microbe Interact. 30, 1009–1019. https://doi.org/10.1094/MPMI-07-17-0176-R

Hellweg, C., Pühler, A., Weidner, S., Puhler, A., Weidner, S., 2009. The time course of the transcriptomic response of Sinorhizobium meliloti 1021 following a shift to acidic pH. BMC Microbiol 9, 37. https://doi.org/1471-2180-9-37 [pii]10.1186/1471-2180-9-37

Janczarek, M., 2015. Exopolysaccharide Production in Rhizobia Is Regulated by Environmental Factors, in: Biological Nitrogen Fixation. John Wiley & Sons, Inc, Hoboken, NJ, USA, pp. 365–380. https://doi.org/10.1002/9781119053095.ch36

Jensen, J.B., Peters, N.K., Bhuvaneswari, T. V., 2002. Redundancy in Periplasmic Binding Protein-Dependent Transport Systems for Trehalose, Sucrose, and Maltose in Sinorhizobium meliloti. J. Bacteriol. 184, 2978–2986. https://doi.org/10.1128/JB.184.11.2978-2986.2002

Kida, Y., Inoue, H., Shimizu, T., Kuwano, K., 2007. Serratia marcescens serralysin induces inflammatory responses through protease-activated receptor 2. Infect. Immun. 75, 164–174. https://doi.org/10.1128/IAI.01239-06

Krol, E., Becker, A., 2011. ppGpp in Sinorhizobium meliloti: biosynthesis in response to sudden nutritional downshifts and modulation of the transcriptome. Mol. Microbiol. 81, 1233–54. https://doi.org/10.1111/j.1365-2958.2011.07752.x

Lambert, A., Osteras, M., Mandon, K., Poggi, M.C., Le Rudulier, D., 2001. Fructose uptake in Sinorhizobium meliloti is mediated by a high-affinity ATP-binding cassette transport system. J Bacteriol 183, 4709–4717. https://doi.org/10.1128/JB.183.16.4709-4717.2001

Lang, C., Barnett, M.J., Fisher, R.F., Smith, L.S., Diodati, M.E., Long, S.R., 2018. Most Sinorhizobium meliloti Extracytoplasmic Function Sigma Factors Control Accessory Functions. mSphere 3. https://doi.org/10.1128/mspheredirect.00454-18

Langridge, G.C., Phan, M.D., Turner, D.J., Perkins, T.T., Parts, L., Haase, J., Charles, I., Maskell, D.J., Peters, S.E., Dougan, G., Wain, J., Parkhill, J., Turner, A.K., 2009. Simultaneous assay of every Salmonella Typhi gene using one million transposon mutants. Genome Res 19, 2308–2316. https://doi.org/10.1101/gr.097097.109

Lipa, P., Vinardell, J.M., Janczarek, M., 2019. Transcriptomic studies reveal that the Rhizobium leguminosarum serine/threonine protein phosphatase PssZ has a role in the synthesis of cell-surface components, nutrient utilization, and other cellular processes. Int. J. Mol. Sci. 20. https://doi.org/10.3390/ijms20122905

Lipa, P., Vinardell, J.M., Kopcińska, J., Zdybicka-Barabas, A., Janczarek, M., 2018. Mutation in the pssZ gene negatively impacts exopolysaccharide synthesis, surface properties, and symbiosis of Rhizobium leguminosarum bv. trifolii with clover. Genes (Basel). 9. https://doi.org/10.3390/genes9070369

Liu, Z., Beskrovnaya, P., Melnyk, R.A., Hossain, S.S., Khorasani, S., O’Sullivan, L.R., Wiesmann, C.L., Bush, J., Richard, J.D., Haney, C.H., 2018. A Genome-Wide Screen Identifies Genes in Rhizosphere-Associated Pseudomonas Required to Evade Plant Defenses. MBio 9. https://doi.org/10.1128/mBio.00433-18

Long, S., McCune, S., Walker, G.C., 1988. Symbiotic loci of Rhizobium meliloti identified by random TnphoA mutagenesis. J. Bacteriol. 170, 4257–4265. https://doi.org/10.1128/jb.170.9.4257-4265.1988

Lynch, D., O’Brien, J., Welch, T., Clarke, P., Cuiv, P.O., Crosa, J.H., O’Connell, M., 2001. Genetic organization of the region encoding regulation, biosynthesis, and transport of rhizobactin 1021, a siderophore produced by Sinorhizobium meliloti. J Bacteriol 183, 2576–2585. https://doi.org/10.1128/JB.183.8.2576-2585.2001

Maier, R.J., Triplett, E.W., 1996. Toward more productive, efficient, and competitive nitrogen-fixing symbiotic bacteria, Critical Reviews in Plant Sciences. https://doi.org/10.1080/07352689609701941

Maki, H., Sekiguchi, M., 1992. MutT protein specifically hydrolyses a potent mutagenic substrate for DNA synthesis. Nature 355, 273–275. https://doi.org/10.1038/355273a0

Materon, L.A., Weaver, R.W., 1985. Inoculant maturity influences survival of rhizobia on seed. Appl. Environ. Microbiol. 49, 465–467.

Miller-Williams, M., Loewen, P.C., Oresnik, I.J., 2006. Isolation of salt-sensitive mutants of Sinorhizobium meliloti strain Rm1021. Microbiology 152, 2049–2059. https://doi.org/152/7/2049 [pii]10.1099/mic.0.28937-0

Oldroyd, G.E.D., 2013. Speak, friend, and enter: signalling systems that promote beneficial symbiotic associations in plants. Nat. Rev. Microbiol. 11, 252–63. https://doi.org/10.1038/nrmicro2990

Olivares, J., Bedmar, E.J., Sanjuán, J., 2013. Biological nitrogen fixation in the context of global change. Mol. Plant. Microbe. Interact. 26, 486–94. https://doi.org/10.1094/MPMI-12-12-0293-CR

Pech-Canul, Á., Nogales, J., Miranda-Molina, A., Álvarez, L., Geiger, O., Soto, M.J., López-Lara, I.M., 2011. FadD Is Required for Utilization of Endogenous Fatty Acids Released from Membrane Lipids. J. Bacteriol. 193, 6295–6304. https://doi.org/10.1128/JB.05450-11

Peoples, M.B., Herridge, D.E., Ladha, J.K., 1995. Biological nitrogen fixation: An efficient source of nitrogen for sustainable agricultural production? Plant Soil 174, 3–28.

Perry, B.J., Akter, M.S., Yost, C.K., 2016a. The Use of Transposon Insertion Sequencing to Interrogate the Core Functional Genome of the Legume Symbiont Rhizobium leguminosarum. Front. Microbiol. 7, 1873. https://doi.org/10.3389/fmicb.2016.01873

Perry, B.J., Wheatley, R.M., Ramachandran, V.K., Poole, P., Yost, C.K., 2016b. High-throughput Transposon Mutagenesis Screening of Pea Symbiont Rhizobium leguminosarum to Investigate Colonization of the Germinating Pea Spermosphere and Radicle, in: 12th European Nitrogen Fixation Conference. BudaPest, Hungary, p. ENFC 2016 Book of abstracts online, page 222, POST.

Pobigaylo, N., Wetter, D., Szymczak, S., Schiller, U., Kurtz, S., Meyer, F., Nattkemper, T.W., Becker, A., 2006. Construction of a large signature-tagged mini-Tn5 transposon library and its application to mutagenesis of Sinorhizobium meliloti. Appl Env. Microbiol 72, 4329–4337. https://doi.org/72/6/4329 [pii] 10.1128/AEM.03072-05

Poole, P., Ramachandran, V., Terpolilli, J., 2018. Rhizobia: From saprophytes to endosymbionts. Nat. Rev. Microbiol. 16, 291–303. https://doi.org/10.1038/nrmicro.2017.171

Price, M.N., Wetmore, K.M., Waters, R.J., Callaghan, M., Ray, J., Liu, H., Kuehl, J. V., Melnyk, R.A., Lamson, J.S., Suh, Y., Carlson, H.K., Esquivel, Z., Sadeeshkumar, H., Chakraborty, R., Zane, G.M., Rubin, B.E., Wall, J.D., Visel, A., Bristow, J., Blow, M.J., Arkin, A.P., Deutschbauer, A.M., 2018. Mutant phenotypes for thousands of bacterial genes of unknown function. Nature 557, 503–509. https://doi.org/10.1038/s41586-018-0124-0

Roughley, R.J., Vincent, J.M., 1967. Growth and Survival of Rhizobium spp. in Peat Culture. J. Appl. Bacteriol. 30, 362–376. https://doi.org/10.1111/j.1365-2672.1967.tb00310.x

Royet, K., Parisot, N., Rodrigue, A., Gueguen, E., Condemine, G., 2019. Identification by Tn-seq of Dickeya dadantii genes required for survival in chicory plants. Mol. Plant Pathol. 20, 287–306. https://doi.org/10.1111/mpp.12754

Rutten, P.J., Poole, P.S., 2019. Oxygen regulatory mechanisms of nitrogen fixation in rhizobia. Adv. Microb. Physiol. 75, 325–389. https://doi.org/10.1016/bs.ampbs.2019.08.001

Salas, M.E., Lozano, M.J., López, J.L., Draghi, W.O., Serrania, J., Torres Tejerizo, G.A., Albicoro, F.J., Nilsson, J.F., Pistorio, M., Del Papa, M.F., Parisi, G., Becker, A., Lagares, A., 2017. Specificity traits consistent with legume-rhizobia coevolution displayed by Ensifer meliloti rhizosphere colonization. Environ. Microbiol. 19, 3423–3438. https://doi.org/10.1111/1462-2920.13820

Santos, M.R., Cosme, A.M., Becker, J.D., Medeiros, J.M.C., Mata, M.F., Moreira, L.M., 2010. Absence of functional TolC protein causes increased stress response gene expression in Sinorhizobium meliloti. BMC Microbiol. 10, 180. https://doi.org/10.1186/1471-2180-10-180

Sengupta, S., Ghosh, S., Nagaraja, V., 2008. Moonlighting function of glutamate racemase from Mycobacterium tuberculosis: Racemization and DNA gyrase inhibition are two independent activities of the enzyme. Microbiology 154, 2796–2803. https://doi.org/10.1099/mic.0.2008/020933-0

Serrania, J., Johner, T., Rupp, O., Goesmann, A., Becker, A., 2017. Massive parallel insertion site sequencing of an arrayed Sinorhizobium meliloti signature-tagged mini-Tn 5 transposon mutant library. J. Biotechnol. 257, 9–12. https://doi.org/10.1016/j.jbiotec.2017.02.019

Silva, C., Kan, F.L., Martínez-Romero, E., 2007. Population genetic structure of Sinorhizobium meliloti and S. medicae isolated from nodules of Medicago spp. in Mexico. FEMS Microbiol. Ecol. 60, 477–489. https://doi.org/10.1111/j.1574-6941.2007.00301.x

Sivakumar, R., Ranjani, J., Vishnu, U.S., Jayashree, S., Lozano, G.L., Miles, J., Broderick, N.A., Guan, C., Gunasekaran, P., Handelsman, J., Rajendhran, J., 2019. Evaluation of InSeq To Identify Genes Essential for Pseudomonas aeruginosa PGPR2 Corn Root Colonization. G3 Genes|Genomes|Genetics g3.200928.2018. https://doi.org/10.1534/g3.118.200928

Smith, R.S., 1992. Legume inoculant formulation and application. Can. J. Microbiol. 38, 485–492. https://doi.org/10.1139/m92-080

Stephens, J.H.G., Rask, H.M., 2000. Inoculant production and formulation. F. Crop. Res. 65, 249–258. https://doi.org/10.1016/S0378-4290(99)00090-8

Sun, J., Nishiyama, T., Shimizu, K., Kadota, K., 2013. TCC: an R package for comparing tag count data with robust normalization strategies. BMC Bioinformatics 14, 219. https://doi.org/10.1186/1471-2105-14-219

Taboada, B., Ciria, R., Martinez-Guerrero, C.E., Merino, E., 2012. ProOpDB: Prokaryotic operon database. Nucleic Acids Res. 40, D627. https://doi.org/10.1093/nar/gkr1020

Tian, C.F.F., Garnerone, A.-M.A.-M., Mathieu-Demazière, C., Masson-Boivin, C., Batut, J., Mathieu-Demaziere, C., Masson-Boivin, C., Batut, J., 2012. Plant-activated bacterial receptor adenylate cyclases modulate epidermal infection in the Sinorhizobium meliloti-Medicago symbiosis. Proc. Natl. Acad. Sci. U. S. A. 109, 6751–6. https://doi.org/10.1073/pnas.1120260109

van Opijnen, T., Bodi, K.L., Camilli, A., 2009. Tn-seq: high-throughput parallel sequencing for fitness and genetic interaction studies in microorganisms. Nat. Methods 6, 767–72. https://doi.org/10.1038/nmeth.1377

van Opijnen, T., Camilli, A., 2013. Transposon insertion sequencing: a new tool for systems-level analysis of microorganisms. Nat. Rev. Microbiol. 11, 435–42. https://doi.org/10.1038/nrmicro3033

Viguier, C., p, O.C., Clarke, P., O’Connell, M., 2005. RirA is the iron response regulator of the rhizobactin 1021 biosynthesis and transport genes in Sinorhizobium meliloti 2011. FEMS Microbiol Lett 246, 235–242. https://doi.org/S0378-1097(05)00234-X [pii]10.1016/j.femsle.2005.04.012

Wetmore, K.M., Price, M.N., Waters, R.J., Lamson, J.S., He, J., Hoover, C. a., Blow, M.J., Bristow, J., Butland, G., Arkin, A.P., Deutschbauer, A., 2015. Rapid Quantification of Mutant Fitness in Diverse Bacteria by Sequencing Randomly Bar-Coded Transposons. MBio 6, e00306–15. https://doi.org/10.1128/mBio.00306-15

Wheatley, R.M., Ford, B.L., Li, L., Aroney, S.T.N., Knights, H.E., Ledermann, R., East, A.K., Ramachandran, V.K., Poole, P.S., 2020. Lifestyle adaptations of Rhizobium from rhizosphere to symbiosis. Proc. Natl. Acad. Sci. U. S. A. 117, 23823–23834. https://doi.org/10.1073/pnas.2009094117

Wheatley, R.M., Ramachandran, V.K., Geddes, B.A., Perry, B.J., Yost, C.K., Poole, P.S., 2017. Role of O 2 in the Growth of Rhizobium leguminosarum bv. viciae 3841 on Glucose and Succinate. J. Bacteriol. 199, e00572–16. https://doi.org/10.1128/JB.00572-16

Zou, L., Gastebois, A., Mathieu-Demazière, C., Sorroche, F., Masson-Boivin, C., Batut, J., Garnerone, A.-M., 2017. Transcriptomic Insight in the Control of Legume Root Secondary Infection by the Sinorhizobium meliloti Transcriptional Regulator Clr. Front. Microbiol. 8, 1–11. https://doi.org/10.3389/fmicb.2017.01236

